# PIP_2_ regulation of TRPC5 channel activation and desensitization

**DOI:** 10.1101/2021.01.25.428089

**Authors:** Mehek Ningoo, Leigh D. Plant, Anna Greka, Diomedes E. Logothetis

## Abstract

Transient receptor potential canonical type 5 (TRPC5) channels are expressed in the brain and kidney, and have been identified as promising therapeutic targets whose selective inhibition can protect against diseases driven by a leaky kidney filter. They are activated by elevated levels of extracellular Ca^2+^ or application of lanthanide ions but also by G protein (G_q/11_) stimulation. Phosphatidylinositol bis-phosphate (PIP_2_) hydrolysis leads to protein kinase C- (PKC-) mediated phosphorylation of TRPC5 channels and desensitization of their activity. Even though PIP_2_ regulation of TRP channels is being widely studied, the roles of PIP_2_ in maintaining TRPC5 channel activity, the PIP_2_ involvement in channel stimulation by its hydrolysis product diacyl glycerol (DAG), or the desensitization of activity by DAG-stimulated PKC activity remain unclear. Here, we show that PIP_2_ controls both the PKC-mediated inhibition of TRPC5 currents as well as the activation by DAG and lanthanides and that it accomplishes this through control of gating rather than channel cell surface density. The mechanistic insights achieved by the present work promise to aid in the development of more selective and precise molecules to block TRPC5 channel activity and illuminate new therapeutic opportunities for targeted therapies for a group of diseases for which there is currently a great unmet need.

---

TRPC5 channels belong to the classical transient receptor potential (TRPC) family of nonselective, calcium permeable cation channels (1). They are widely expressed in many tissues, including the brain, where they are involved in fear-related behavior, regulating hippocampal neurite length as well as growth cone morphology, and the kidney, where they are largely implicated in chronic kidney disease. In kidney podocytes, cells essential for the kidney filter, TRPC5 channels may be promising therapeutic targets, because their selective inhibition is protective against diseases driven by a leaky kidney filter in rodents (2–6). Mammalian TRPC5 channels are transiently stimulated by the action of phospholipase C (PLC) enzymes, either GTP-binding protein coupled receptors (GPCRs) coupled to Gq_/11_ that signal through PLCβ_1_ or by tyrosine kinase-coupled receptors through PLCγ_2_ (1–4). Activation of PLC causes the hydrolysis of plasma membrane phosphatidylinositol (4,7) bis-phosphate (PIP_2_) to form inositol 1,4,5-triphosphate (IP_3_) that releases Ca^2+^ from intracellular stores and diacylglycerol (DAG) which activates protein kinase C (PKC) (1–7).

Since the mid-1990s PI(4,5)P_2_ (PIP_2_) has been appreciated to act beyond its role as a precursor to the ubiquitous signaling of its products (e.g. IP_3_, DAG, PIP_3_) as a direct regulator of membrane protein function, especially ion channel proteins (8,9). Co-crystal structures of Kir channels with PIP_2_ have revealed at atomic resolution specific residue interactions with the phosphates at positions 4’ and 5’ of the inositol ring of PI(4,5)P_2_ around the two channel gates of Kir channels (10–12). Microsecond-long MD simulations have revealed how gating molecules, like the Gβγ subunits or Na^+^ ions, open one, the other or both gates of Kir3 channels by enhancing specific residue interactions with PIP_2_ (13). The relationship of TRPC5 with PIP_2_ has not yet been structurally elucidated (14).

TRPC5 channels can also be steadily activated by elevated levels of extracellular Ca^2+^ or application of lanthanide ions, such as lanthanum (La^3+^) and gadolinium (Gd^3+^) (15, 16). These cations bind to an extracellular cation binding site (eCBS) located in the vicinity of the channel’s pore entrance (15, 16). The eCBS consists of two acidic Glu residues, E543 and E595, mutation of either of which to Gln renders TRPC5 lanthanide-insensitive. The Gln mutant channels can still be activated through PLC stimulation, suggesting that the two mechanisms of channel activation are independent (17).

Although it had been thought that the PIP_2_ hydrolysis products IP_3_ and DAG do not themselves activate TRPC5 channels, evidence for a mechanism that renders the channel sensitive to DAG-mediated activation has been presented (2). This mechanism requires Gq-receptor mediated activation of PLC and hydrolysis of PIP_2_ causing a conformational change in the C-terminus of TRPC5 that leads to dissociation of the Na^+^ /H^+^ exchanger regulatory factor (NHERF) protein from a PDZ-binding motif (2). NHERF proteins serve to link integral membrane proteins to the cytoskeleton and NHERF dissociation allows the channel to be activated by DAG (2).

TRPC5 channels that are dependent on PlC*β* or PLC*γ* activation, exhibit current desensitization by a mechanism attributed to PKC-mediated phosphorylation (17,18). In this scheme, DAG-mediated activation of TRPC5 channels precedes activation of PKC which phosphorylates the channel at T972 to cause desensitization by an unknown mechanism (18). PKC phosphorylation of the channel at T972 has been proposed to increase the affinity of the C-terminus of the channel to bind to NHERF1, blocking activation by DAG (2). Mutating the Thr residue to Ala (T972A) protects the channels from being phosphorylated by PKC and prevents desensitization (2), favoring DAG over NHERF binding to result in channel activation. For G protein gated Kir3 channels, PKC-mediated channel phosphorylation has been shown to weaken channel-PIP_2_ interactions and, together with PIP_2_ hydrolysis, to underlie current desensitization (20, 21). To date, even though PIP_2_ regulation of TRP channels is being widely studied (18, 19, 22), the roles of PIP_2_ in maintaining TRPC5 channel activity, its involvement in the DAG-mediated stimulation and in PKC-mediated desensitization of activity remain unclear.

The present work provides evidence that trivalent cation and PLC-mediated activation of TRPC5 allosterically and independently converge on PIP_2_ to gate the channel. Both the PKC-mediated phosphorylation of T972 and direct PIP_2_ depletion contribute to TRPC5 current inhibition. PKC-phosphorylation of the channel weakens channel-PIP_2_ interactions, while DAG-mediated activation of the PKC-insensitive T972A mutant strengthens channel-PIP_2_ interactions. Our findings support a paradigm whereby PIP_2_ hydrolysis and DAG generation play a dual role in G_q/11_ protein-mediated TRPC5 channel activation. First, in the absence of phosphorylation at T972, DAG is able to strengthen channel-PIP_2_ interactions and stimulate activity. Second, DAG activates PKC that in turn phosphorylates the channel at T972, driving activity toward full inhibition by weakening channel-PIP_2_ interactions and enabling loss of PIP_2_ from the channel, given the surrounding depleted levels.

## Results

### Wortmannin speeds Gq-mediated inhibition kinetics of TRPC5 channels

Activation of Gq-signaling evokes a transient TRPC5 current that subsequently decreases in magnitude via a mechanism that is proposed to require phosphorylation of the channel at threonine 972 by PKC (18). To determine if the PKC-mediated decrease in TRPC5 currents is PIP_2_ sensitive, we used whole-cell patch-clamp recording to study HEK293T cells transiently expressing mTRPC5-GFP under experimental conditions where PIP_2_ levels were elevated or lowered. Double-rectifying TRPC5 currents were elicited by voltage ramps from −100 mV to +100 mV following application of 100 μM carbachol (CCh), an agonist at the Gq-coupled muscarinic type-3 (M3) receptors that are endogenously expressed in HEK293T cells (**Fig. 1A**) (24). Following the initial activation of the channel, TRPC5 currents spontaneously decreased in magnitude, as expected. To probe if the rate of PKC-mediated channel inhibition is PIP_2_-sensitive, we increased the level of PIP_2_ in the HEK293 cells through two types of manipulations: First, we increased production of PIP_2_ by co-transfecting mTRPC5-GFP with phosphatidylinositol 4-phosphate 5 kinase (PIP-5K), the enzyme that phosphorylates PI(4)P at position 5’ of the inositol ring. Second, we increased PIP_2_ levels directly by including 200 μM dioctanoyl-glycerol-PIP_2_ (diC_8_-PIP_2_), a soluble analog of PIP_2_ in the patch pipette, as before (25). Both PIP-5K overexpression and inclusion of diC_8_-PIP_2_ in the patch pipette reduced the extent to which TRPC5 currents decreased following activation by CCh (**Fig 1B**). Next, we studied the effect of depleting intracellular PIP_2_ levels by incubating the cells in 20 μM Wortmannin, a fungal sterol that at micromolar concentrations inhibits PI3K and PI4K, of which PI4K is required to generate PIP_2_ (5). Following a one-hour incubation with Wortmannin, CCh-activated TRPC5 currents showed a higher rate of inhibition than untreated controls (**Fig. 1C**). Together, these findings indicate an inverse relationship between PIP_2_ levels and PKC-induced inhibition of channel currents.

**Figure 1.**
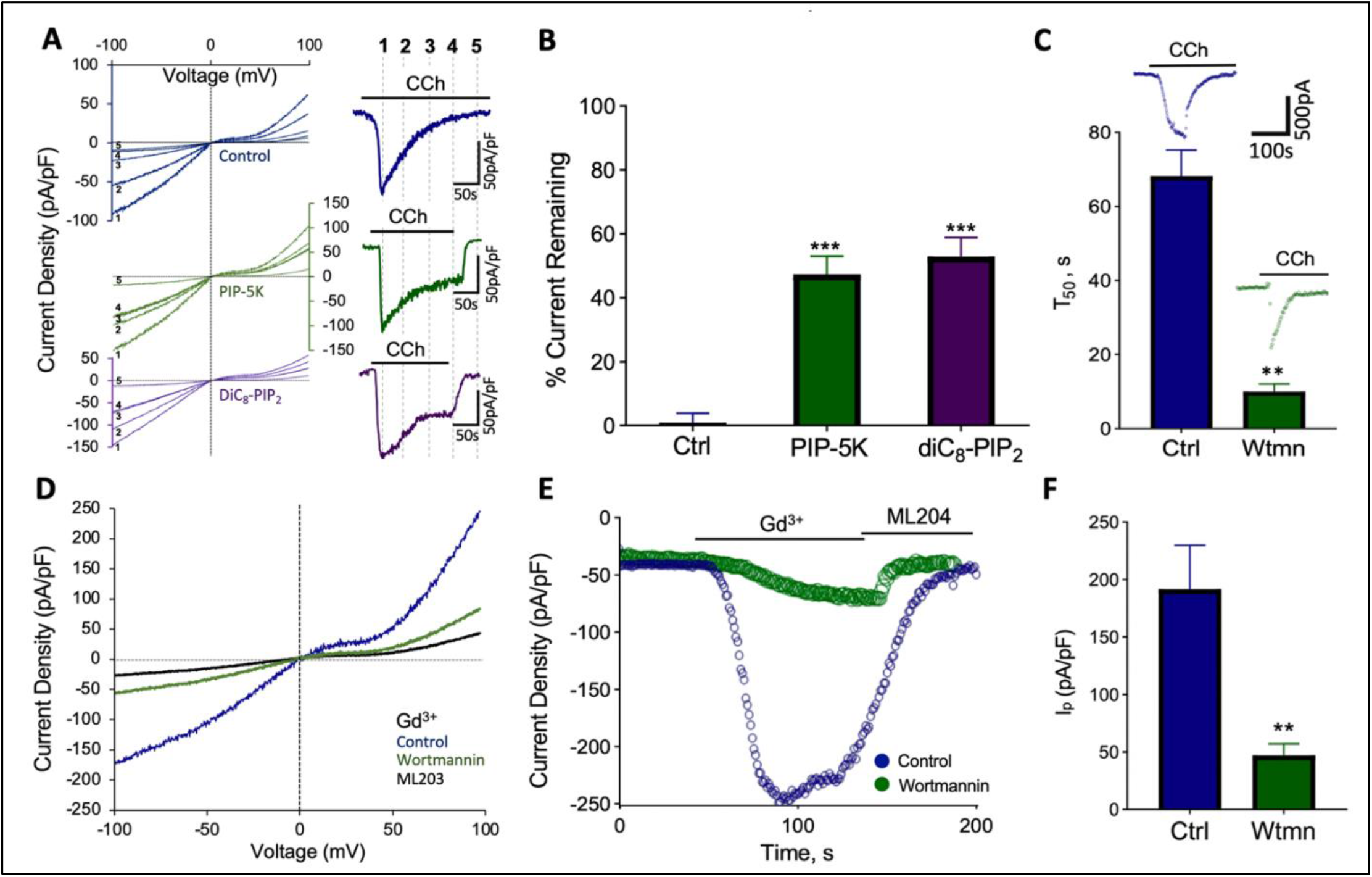
PIP_2_ implicated in PKC-mediated desensitization and promotion of Gd^3+^-activated TRPC5 currents. **A**) Left: Example current-density, voltage relationships for HEK293T cells expression mTRPC5-GFP. Currents were evoked by a ramp from −100 mV to 100 mV follow application of 100 μM CCh. Cells were studied under control conditions (blue), or with PIP-5K co-expression (green) or with 200 μM diC_8_-PIP_2_ in the pipette (purple). The spontaneous decrease in current is illustrated by sweeps labeled 1-5, which correspond to 50 s intervals, as illustrated on the exemplar time courses (Right) **B**) Mean of % current remaining during 100 μM CCh treatment in control HEK293T cells expressing mTRPC5-GFP (n=12, mean 0.89 ± 0.85), with overexpression of PIP-5K (n=9, mean 47.07 ± 1.81) and with diC_8_-PIP_2_ in the pipette (n=10, mean 52.32 ± 1.86 **C**) Bar graph of time taken from peak to 50% current decay (T_50_) of control HEK293T cells expressing mTRPC5-GFP (n=11, mean 68.82 ± 7.52) and after treatment with 20 μM wortmannin for 1 hour (n=9, mean 10.23 ± 1.91); Top: Representative whole-cell patch-clamp recording of HEK293T cells expressing TRPC5-GFP activated by 100 μM CCh. **D**) Current density voltage curves of the ± 100mV ramp of 100 μM Gd^3+^ activation in TRPC5-GFP expressing HEK293T cells (control) and after 1 hour treatment with 20 μM Wortmannin. **E**) Representative whole-cell Current density (pA/pF) curves observed in HEK293T cells overexpressing mTRPC5-GFP activated with 100 μM GdCl_3_ and upon 20 μM wortmannin treatment for 1 hour. **F**) Bar graph of Ip (peak current density-pA/pF) of control HEK293T cells expressing TRPC5-GFP (n=5, mean 191 ± 45.09) and cells treated with wortmannin (n=5, mean 49.35 ± 9.94). P-values established using Student’s t-test. \*\**p<0.001*, \*\*\**p<0.0001*

TRPC5 channels are also activated by extracellular trivalent lanthanide ions via a mechanism that is independent of Gq-signaling, does not activate PKC-mediated phosphorylation, and is not associated with a subsequent decrease in current (25). Application of 100 μM Gd^3+^ activated doubly-rectifying currents that were blocked by the TRPC5 inhibitor ML204 but did not decrease in magnitude over time (**Fig. 1D**) (26). Incubation with 20 μM Wortmannin decreased the peak Gd^3+^-elicited currents by 74.16 ± 0.75% compared to untreated control cells (**Fig. 1E, F**). Taken together, the results of figure 1 implicate multiple roles for PIP_2_ in the activation and inhibition of TRPC5 channels by both independent gating mechanisms, stimulation by Gq signaling and lanthanides.

### PMA-mediated inhibition weakens channel-PIP_2_ interactions

The spontaneous decrease in TRPC5 current observed following activation by CCh is proposed to be mediated by subsequent PKC-mediated phosphorylation of the channel (18). The two key molecules of interest in the desensitization pathway are the PKC enzymes which induce desensitization by phosphorylation and PIP_2_, which opposes desensitization. To control the kinetics of TRPC5 current inhibition, we used optogenetic-activation of 5’-phosphatase to dephosphorylate PIP_2_ in cells illuminated by 460 nm (blue) light. Briefly, blue light-induces dimerization between two plant proteins, cryptochrome 2 (CRY2) and the transcription factor CIBN to control the plasma membrane PIP_2_ levels rapidly, locally, and reversibly. The 5’-phosphatase domain of OCRL (5’-ptaseOCRL), which acts on PI(4,5)P_2_ and PI(3,4,5)P_3_, is fused to the photolyase homology region domain of CRY2 (23). Stimulation of the CRY2-binding domain, CIBN, results in nearly instantaneous recruitment of 5’-ptaseOCRL to the plasma membrane, causing rapid PI(4,5)P_2_ dephosphorylation to PI(4)P (23). We expressed the TRPC5 channel, light-activated CRY2-5’PTASE_OCRL_ and CIBN-CAAX-GFP proteins in HEK-293T cells in order to perform the blue-light activated phosphatase experiment in the whole-cell mode of the patch-clamp technique. Gd^3+^-activated inward currents were allowed to stabilize before a blue light (460 nm, shown by the blue panels) was shone to activate the inositol 5’-phosphatase and deplete membrane PIP_2_ until the current declined and reached a steady state (**Fig. 2A**).

**Figure 2.**
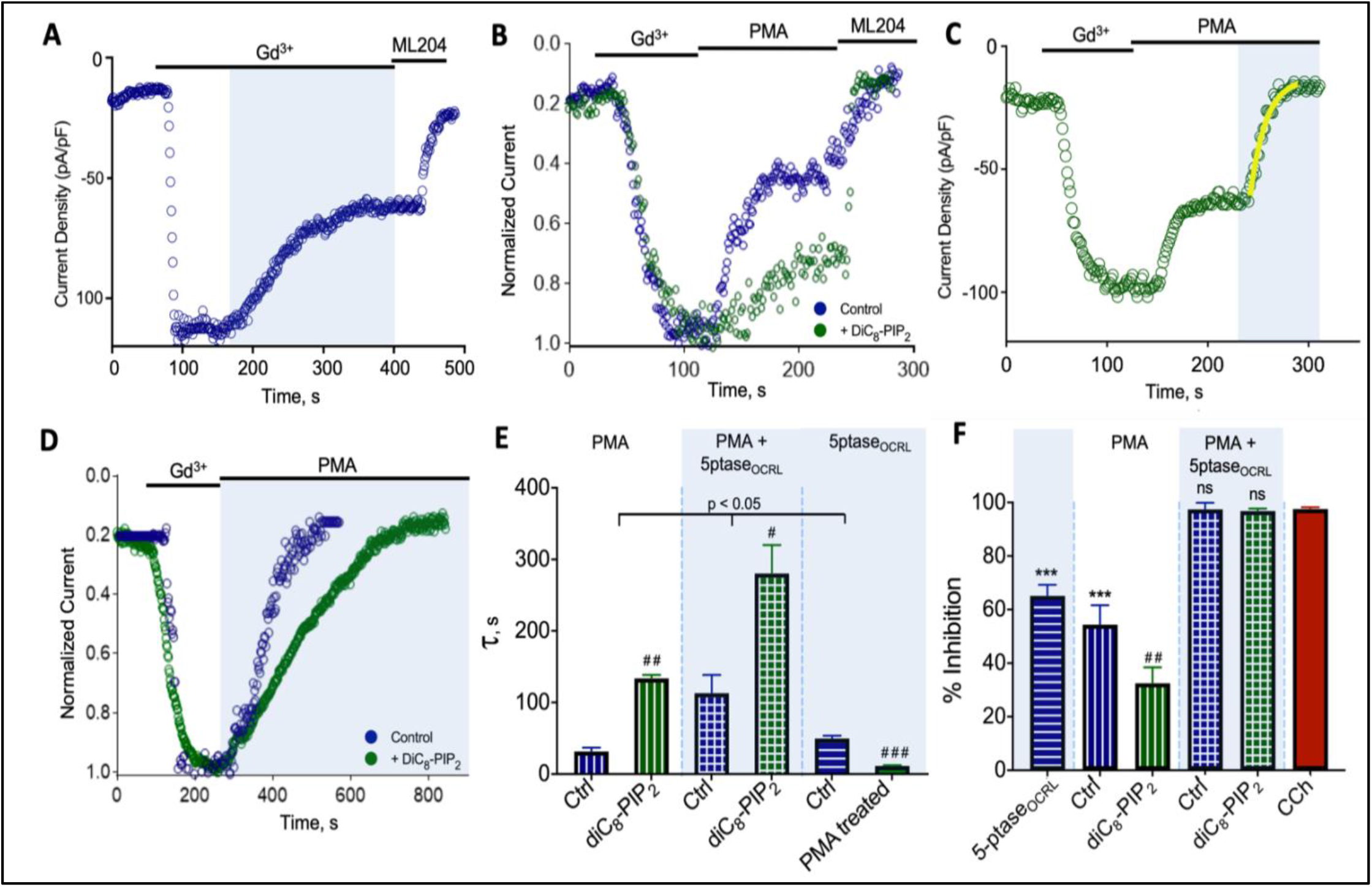
TRPC5 current inhibition by PKC-mediated phosphorylation and/or PIP_2_ dephosphorylation reveals an underlying decrease in channel-PIP_2_ interactions. **A**) Whole-cell patch clamp recording of HEK293T cells expressing TRPC5-GFP, light-activated CRY2-5’PTASE_OCRL_ and CIBN-CAAX-GFP (see Methods), inward current activated by 100 *μ*M GdCl_3_ with channel current decrease in response to light-activated metabolism of PIP_2_ and remaining current blocked by 3 *μ*M ML204. **B**) Inhibition observed by PKC-activator PMA without/with 200 μM diC_8_-PIP_2_ in the pipette. **C**) HEK-293T cells expressing TRPC5-GFP, CRY2-5’ptase and CIBN-CAAX-GFP were activated using 100 μM GdCl_3_, 200 nM PMA was applied to activate PKC enzymes followed by blue-light exposure. **D**) Inhibition observed by simultaneous application of PKC-activator PMA and activation of light-activated inositol phosphatase without/with 200 μM diC_8_-PIP_2_ in the pipette. **E**) Bar graph of the mean decay constant of PMA-mediated inhibition alone (n=5, 31.44 ± 5.55) and with diC_8_-PIP_2_ (n=5, 133.95± 4.48), simultaneous PMA and 5’-ptase_OCRL_ mediated inhibition (n=6, 112.97 ± 25.67) and with diC_8_-PIP_2_ (n=5, 280.33 ± 39.77), and 5’-ptase_OCRL_ mediated inhibition alone (n=8, 52.57 ± 4.70) and after PMA treatment (n=5, 11.43 ± 0.92). **F**) Bar graph summary of mean % current inhibition (* values, compared with CCh) by 5’-ptase_OCRL_ (n=8, 65.18 ± 2.37), PMA mediated inhibition alone (n=5, 54.4 ± 4.17) and with diC_8_-PIP_2_ (n=5, 32.47± 3.48), simultaneous PMA and 5’-ptase_OCRL_ mediated inhibition (n=6, 97.51 ± 1.38) and with diC_8_-PIP_2_ (n=5, 96.87 ± 0.47), and when activated using 100μM CCh (n=12, 97.61 ± 0.33). P-values established using Students’ t-test. #; denotes comparison with experimental Ctrl, #*p<0.01,* ## *p<0.001*, ###*p<0.0001*

Next, to examine the effect of PKC-mediated inhibition of the channel without causing a concurrent hydrolysis of PIP_2_, PKC was activated using 100nM phorbol-12-myristate-13-acetate (PMA) (**Fig. 2B**). To assess the role of PIP_2_ on the PKC effect, we increased intracellular PIP_2_ levels using diC_8_-PIP_2_ in the pipette (**Fig. 2B**). We observed that inclusion of diC_8_-PIP_2_ in the pipette solution decreased the rate and extent of desensitization induced by PMA (**Fig. 2E, F**). This emphasizes the significance of channel-PIP_2_ interaction strength on the inhibition of channel by PKC-mediated phosphorylation. We observed that selective dephosphorylation of PIP_2_ (blue-light phosphatase assay) and activation of PKC enzymes using the drug PMA, individually inhibit approximately 50% of the channel current (**Fig. 2A, B**). Next, we probed the effect of PKC mediated effects on the strength of channel-PIP_2_ interaction by comparing the kinetics of inhibition due to dephosphorylation of PIP_2_ before and after the channel was treated with PMA. After the PMA-induced inhibition of TRPC5 currents reached a steady state (**Fig. 2C**), blue-light illumination yielded faster inhibition kinetics, suggesting that the PKC-mediated phosphorylation decreases channel-PIP_2_ interactions (**Fig. 2A, C, E**).

To test whether the complete current inhibition observed during PLC-activation (see **Fig. 1A**) could be mimicked by simultaneous PIP_2_ hydrolysis and PKC-mediated effects, the Gd^3+^-activated channel was simultaneously treated with the PKC-activator PMA and blue-light to cause dephosphorylation of PIP_2_. This resulted in a complete inhibition (**Fig. 2D, F**) of channel currents similar to that observed in the Gq-activated system (**Fig. 1A**), suggesting that PKC activation and PIP_2_ hydrolysis are both involved to cause complete channel inhibition when TRPC5 channels are activated by the Gq-receptor signaling pathway. Next, we determined the effect of increased levels of intracellular PIP_2_ by repeating the same experiment in cells studied with 200μM diC_8_-PIP_2_ in the recording pipette. Elevated levels of PIP_2_, and thus increased channel-PIP_2_ interactions, decreased the rate of channel inhibition by simultaneous PIP_2_ dephosphorylation and PKC activation (**Fig. 2D, E**) but did not affect the extent of inhibition (**Fig. 2D, F**). Given these results, we conclude that the rate of desensitization of TRPC5 channels by PKC is inversely affected by intracellular PIP_2_ levels.

### OAG strengthens TRPC5 channel-PIP_2_ interactions to stimulate channel activity

The T972A TRPC5 mutant is PKC-insensitive, abolishing the desensitization observed on Gq-mediated activation of the channel and enabling direct activation by OAG (2, 17). Since our experiments thus far suggested that PKC mediated phosphorylation is dependent on channel-PIP_2_ interactions, we investigated further whether the PKC-insensitive mutant channel differs from the wild-type channel in its interaction with PIP_2_. We used 100 μM GdCl_3_ to activate the wild-type and T972A mutant channels to bypass the activation of the PLC pathway and assess channel-PIP_2_ strength by exposure to blue-light (**Fig. 3A**). The mutant channel exhibited a significantly slower inhibition (**Fig. 3A, B**) than the wild-type channel indicating a stronger channel-PIP_2_ interaction compared to the wild-type. This result also suggested the occurrence of basal PKC-dependent phosphorylation under unstimulated conditions. From this observation we can conclude that the PKC-insensitive T972A mutant channel has stronger channel-PIP_2_ interactions compared to the wild-type channel. The TRPC5 channel is thought to be activated by DAG, which is endogenously produced through hydrolysis of PIP_2_ by PLC enzymes. To understand the mechanism by which DAG is activating the channel and whether it occurs through modulating the channel’s interaction with the remaining non-hydrolyzed PIP_2_, we used a saturating concentration of OAG (200μM) to activate the channel and compared its kinetics of inhibition upon blue-light induced PIP_2_ dephosphorylation with that of a saturating concentration of Gd^3+^ (150μM). Channels activated by OAG showed a slower inhibition upon dephosphorylation of PIP_2_ than those activated by Gd^3+^ (**Fig. 3D**) indicating that OAG activation is characterized by stronger interactions of the channel with PIP_2_. To look into the effect of PIP_2_ levels on OAG-activated currents, we depleted PIP_2_ levels using Wortmannin and increased intracellular PIP_2_ levels by including diC_8_-PIP_2_ in the pipette solution We observed that upon Wortmannin incubation, the peak current density of OAG-activated currents was significantly smaller than control (data not shown) or 200 μM PIP_2_ in the pipette solution. Interestingly, these varied PIP_2_ levels that affect channel-PIP_2_ interaction strength also affected the rate of inhibition of OAG-activated currents upon dephosphorylation of the membrane PIP_2_ using the blue-light activated phosphatase assay. This indicated that the activation of the channel by DAG, similar to channel desensitization by PKC (**Fig. 2**), is dependent on the channel-PIP_2_ interaction strength.

**Figure 3.**
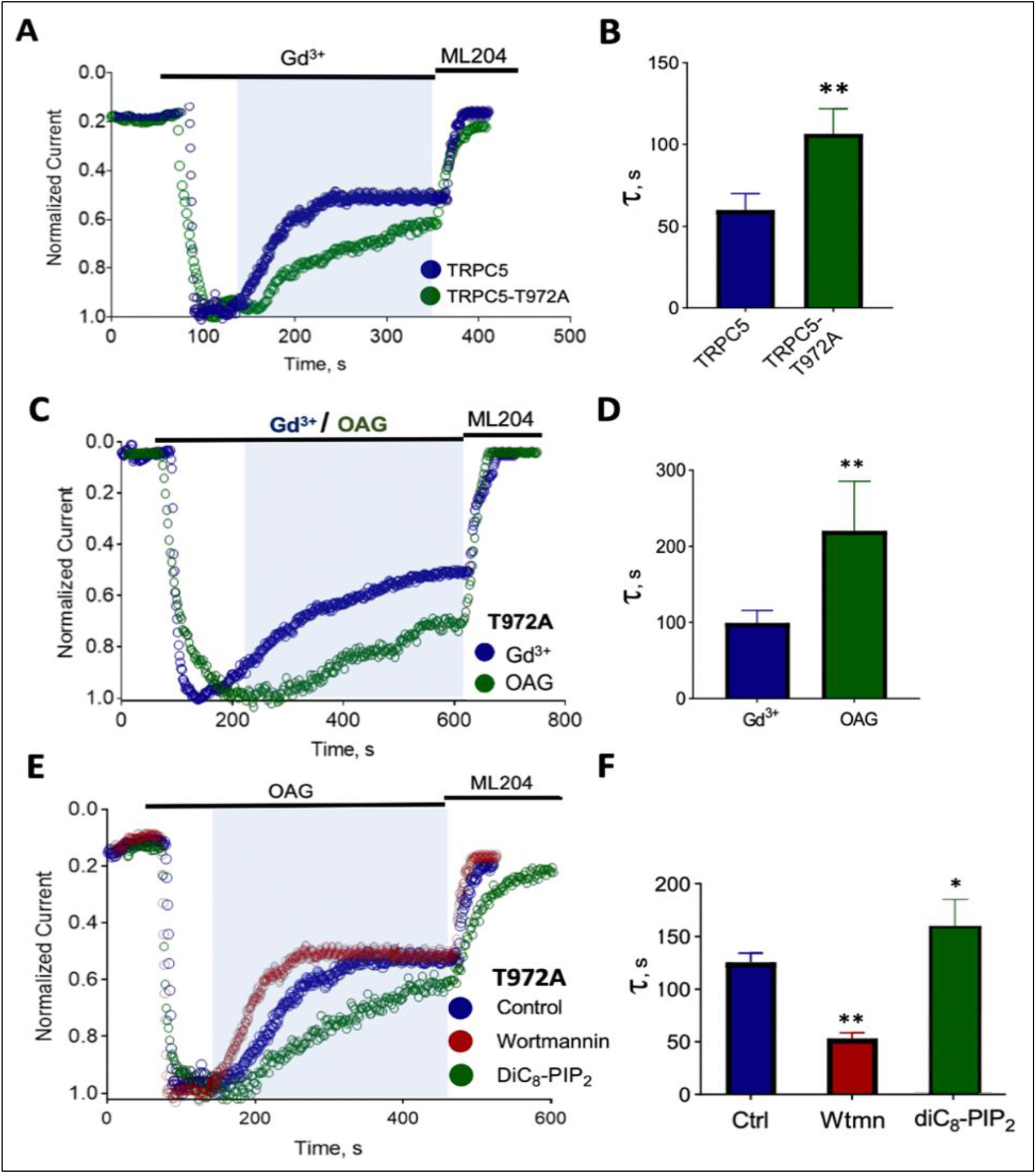
OAG mediated activation of TRPC5 channels shows enhanced channel-PIP_2_ interaction strength. **A)** HEK-293T cells expressing TRPC5-GFP/TRPC5-T972A-GFP, CRY2-5’ptase and CIBN-CAAX-GFP were activated using 100 μM GdCl3 and effect of blue-light exposure was observed. **B)** Bar graph of the mean decay constant of inhibition for TRPC5 (n=6, 60 ± 5.77) for mTRPC5-T972A (n=4, 106.67 ± 8.82.) **C**) HEK-293T cells expressing TRPC5-T972A-GFP, CRY2-5’ptase and CIBN-CAAX-GFP were activated using saturated concentration of Gd^3+^ (150 μM) or OAG (200 μM) and effect of blue-light exposure was observed. **D**) Bar graph of the mean decay constant of inhibition when activated by 150μM Gd^3+^ (n=5, 99.55 ± 8.15) and 200μM of OAG (n=5, 220.5 ± 32.34). **E**) HEK-293T cells expressing TRPC5-T972A-GFP, CRY2-5’ptase and CIBN-CAAX-GFP were activated using 100μM OAG (control), incubated in 20 μM Wortmannin for 1 hour and with 200 μM diC_8_-PIP_2_ in the pipette. **D)** Bar graph of the mean decay constant of inhibition for control (n=5, 125.5 ± 4.35), with wortmannin (n=5, 53.25 ± 2.69), and with diC_8_ PIP_2_ (n=5, 178.25 ± 5.45). P-values established using Students’ t-test. \**p<0.01*, \*\**p<0.001*, \*\*\**p<0.0001*

### The surface-density of TRPC5 channels is not regulated by PIP_2_ or OAG

YFP-tagged TRPC5-T972A channels also expressed readily in HEK293T cells at levels similar to wild type channels (25 ± 1 particles per 10 μm^2^; n = 12) in both intact cells, or when cells were studied in TIRF-patch mode with control solution, 200 μM OAG, or 200 μM diC_8_-PIP_2_ included in the pipette (**Fig. 4C, D**). Like wild type YFP-TRPC5 channels, the number of YFP-TRPC5-T972A channels at the cell membrane was not altered by treating cells with staurosporine, or via activation of 5’-ptase_OCRL_, irrespective of the solution in the patch-pipette (**Fig. 4D**). Together, these findings suggest that changes in the current density of TRPC5 channels observed following changes in the level of PIP_2_, or after activation of PKC result from the regulation of channel activity and not from modified trafficking of channels to, or from the cell membrane.

**Figure 4.**
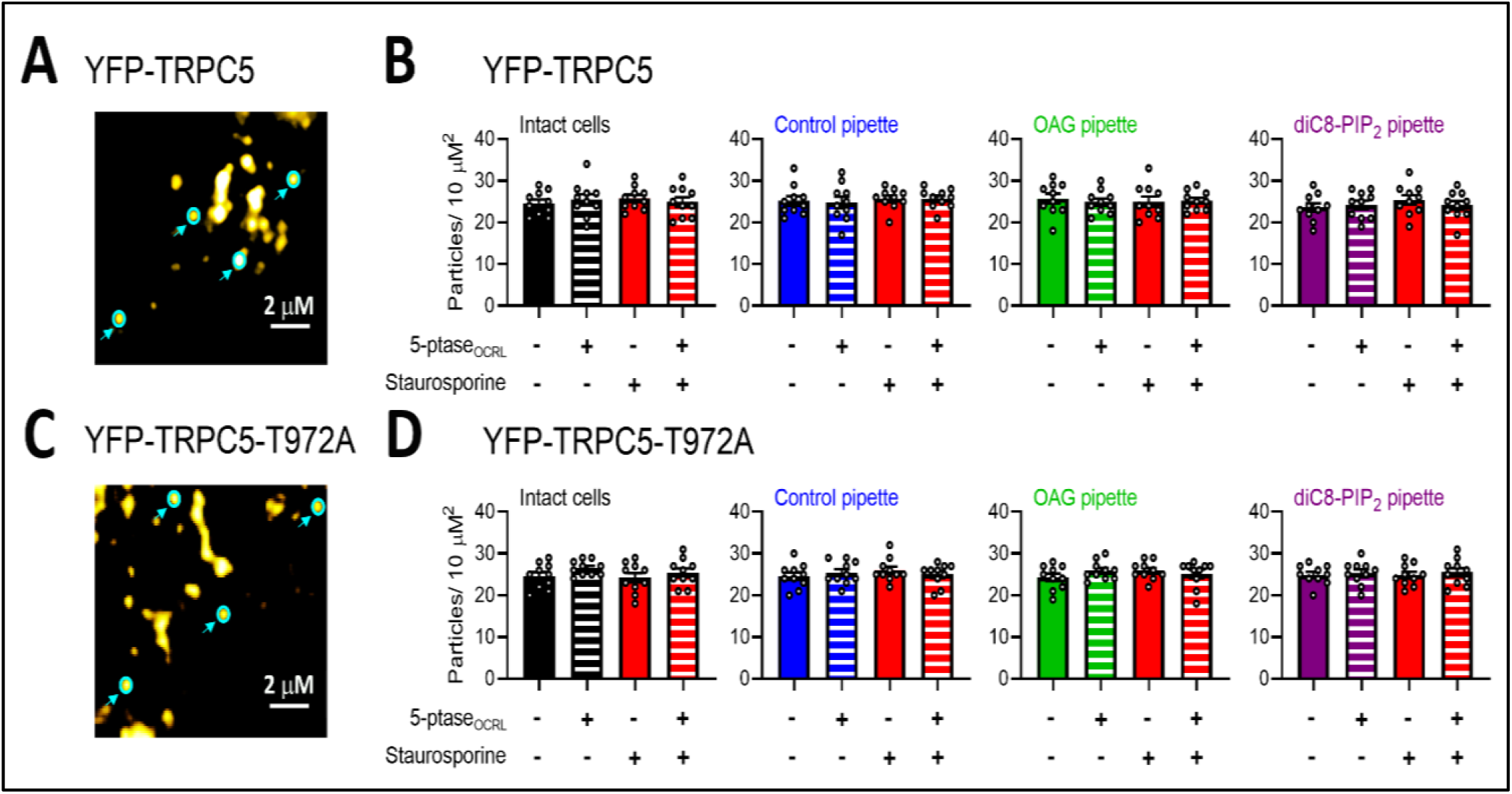
Regulation by OAG or PIP_2_ does not alter the surface-density of YFP-TRPC5 channels. YFP-tagged mTRPC5 or mTRPC5-T972A channels were expressed in HEK293T cells and studied by patch-clamp TIRF. The number of fluorescent particles was determined 200 s after whole-cell mode was established to allow dialysis of the cells with control solution (blue), 200 μM OAG (green) or 200 μM diC8-PIP_2_ (purple). Cells were studied with or without optogenetic activation of 5’-ptaseOCRL (white stripped bars) or following incubation with 1 μM staurosporine (red). Bar graphs represent particle-density as mean ± s.e.m. number of fluorescent particles in the TIRF field in 3-6 random 10 x 10 μm squares per cell and from 4-6 cells per group. **A**) TIRF image showing YFP-tagged wild type mTRPC5 channels at the cell surface. Four example particles corresponding to single TRPC5 channels are highlighted in cyan. **B**) Bar graphs summarizing the density of fluorescent particles indicating no change from control values under any of the conditions studied. **C**) TIRF image showing YFP-tagged mTRPC5-T972A channels at the cell surface. Four example particles corresponding to single TRPC5 channels are highlighted in cyan. **D**) Bar graphs summarizing the density of fluorescent particles indicating no change from control values under any of the conditions studied.

### PIP_2_ prevents PKC-mediated desensitization and promotes OAG-mediated activation in endogenously expressed TRPC5 channels

Since TRPC5 channels are known to be highly expressed in the hippocampus, we utilized the hippocampal neuronal cell line HT-22 to study the channel in a native system (2). TRPC5 channel current was observed upon Gd^3+^application (**Fig. 5A, B**) and like in the HEK293T cell overexpression system it was OAG insensitive (data not shown). To investigate the role of PIP_2_ on PKC-mediated desensitization, we examined the rate and extent of inhibition via PMA with and without diC_8_-PIP_2_ in the patch pipette. In the presence of increased intracellular PIP_2_, PKC-mediated inhibition was slower and less efficient (**Fig. 5C, D**), similar to observations in the HEK293T overexpression system (**Fig. 2E**). Next, to confirm the dependency of DAG-mediated activation on PIP_2_ levels, we first incubated the cells in 1 μM staurosporine to inhibit PKC. This made the channel sensitive to OAG activation, and produced further current stimulation by increased intracellular PIP_2_ levels (**Fig. 5E, F, G**). Having diC_8_-PIP_2_ in the patch-pipette gave higher peak currents compared to control, indicating that PIP_2_ contributes to higher OAG-mediated channel activity. Altogether, these results indicate that PIP_2_ regulation of endogenous TRPC5 channels mirrors its effect in heterologously expressed channels.

**Figure 5.**
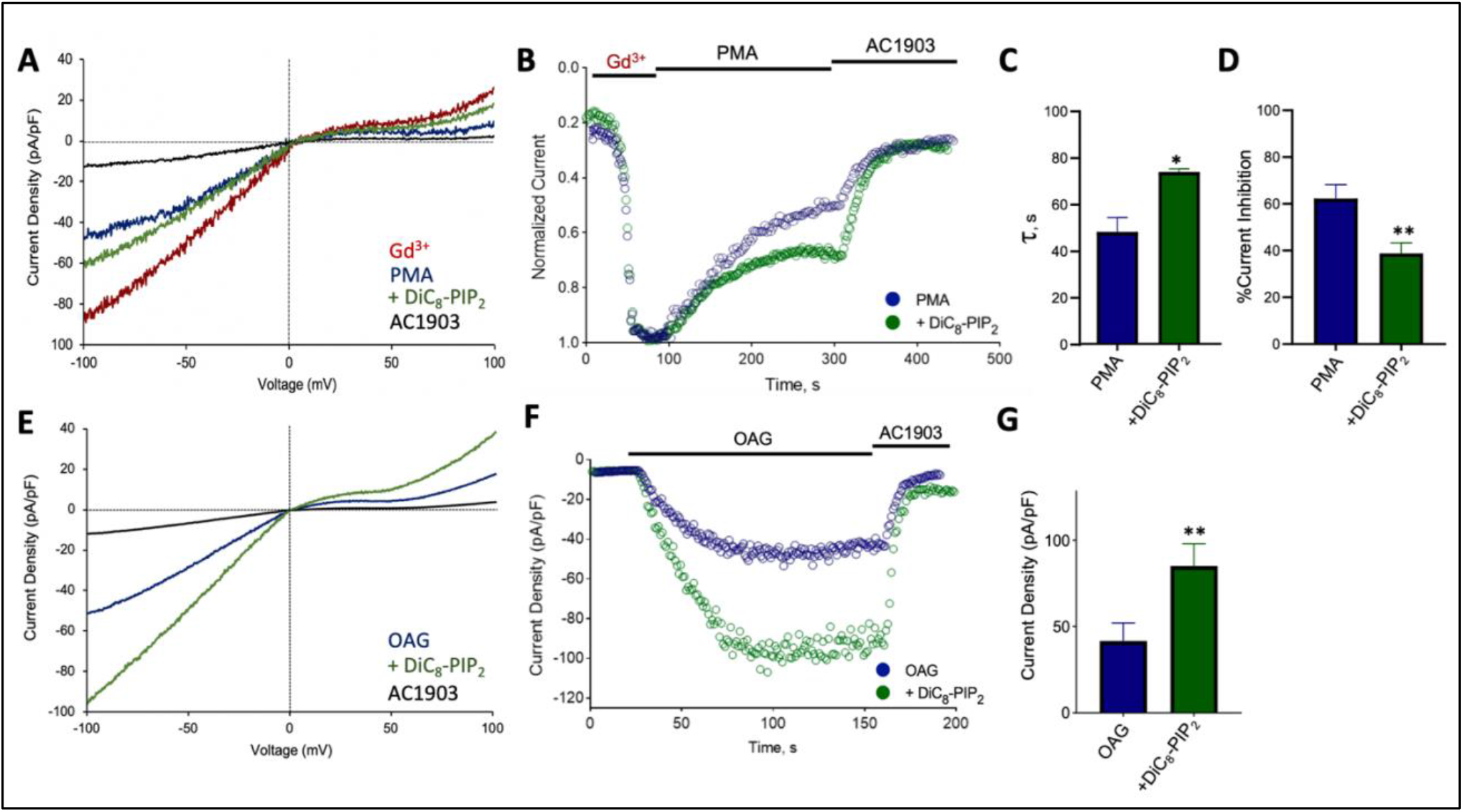
PIP_2_ prevents PKC-mediated desensitization and promotes OAG-mediated activation in endogenously expressed TRPC5 channels. **A**) Current density voltage curves of the ± 100mV ramp of 100 μM Gd^3+^, PMA inhibition with/ without 200 μM diC_8_-PIP_2_ in the pipette and inhibition with 100 μM AC1903. **B**) Representative whole-cell recording of PMA mediated inhibition of Gd^3+^ with/ without 200 μM diC_8_-PIP_2_. **C**) Bar graph summary of mean decay constant of inhibition observed with PMA (Ctrl n=3, 49.5 ± 4.5) and with 200 μM diC_8_-PIP_2_ (n=3, 76 ± 1.3). **D**) Bar graph summary of % current inhibited with PMA (Ctrl n=3, 62.26 ± 2.96) and with 200μM diC_8_-PIP_2_ (n=3, 38.8 ± 3.2). **E**) Current density voltage curves of the ± 100mV ramp in HT-22 cells treated with 1μM staurosporine for 30 mins, of 100 μM OAG activation with/ without 200 μM diC_8_-PIP_2_ in the pipette and inhibition with 100 μM AC1903. **F**) Representative whole-cell recording of 100μM OAG activated currents in HT-22 cells treated with 1μM staurosporine for 30 mins, with/ without 200 μM diC_8_-PIP_2_. **G**) Bar graph summary of peak current density observed with 100 μM OAG (Ctrl n=3, 42.5 ± 11.8) and with 200 μM diC_8_-PIP_2_ (n=3, 81.6 ± 18.3).

## Discussion

Until now, the role of PIP_2_ in the mechanism that regulates TRPC5 channel activity after stimulation of G_q/11_-coupled receptors has remained largely elusive. Even though there is evidence that TRPC5 channels become DAG sensitive upon PLC-mediated hydrolysis of PIP_2_, the specific role of PIP_2_ in channel activation or inhibition had not been probed (2). In this study, we show that TRPC5 channels are functionally coupled to PIP_2_ and that DAG activation as well as PKC-mediated inhibition of the channel, through phosphorylation at T972, involve modulation of channel-PIP_2_ interactions. Our findings consolidate and synthesize prior seemingly discrepant results on the role of PIP_2_ into a single, coherent model.

We found that in trivalent ion-mediated channel gating, Gd^3+^ strengthens channel-PIP_2_ interaction to cause sub-maximal channel activation (see model in **Fig. 6**, strength 4/5). PKC-mediated phosphorylation of the channel (PMA treatment) weakens channel-PIP_2_ interactions and causes partial inhibition (strength 3/5). Similarly, PIP_2_ depletion (5’-phosphatase) inhibits activity partially due to the high-affinity of the channel for PIP_2_ that protects the channel from losing all of its interacting PIP_2_ molecules in the Gd^3+^ activated state (7). A combination of PIP_2_ depletion and PKC-mediated channel phosphorylation fully inhibits currents below basal levels, as the channel is now less able to hold on to its interacting PIP_2_ molecules (strength 1/5). Trivalent ion gating of TRPC5 channels is more straight-forward than Gq-mediated gating as activation and inhibition can be controlled separately as well as simultaneously.

**Figure 6.**
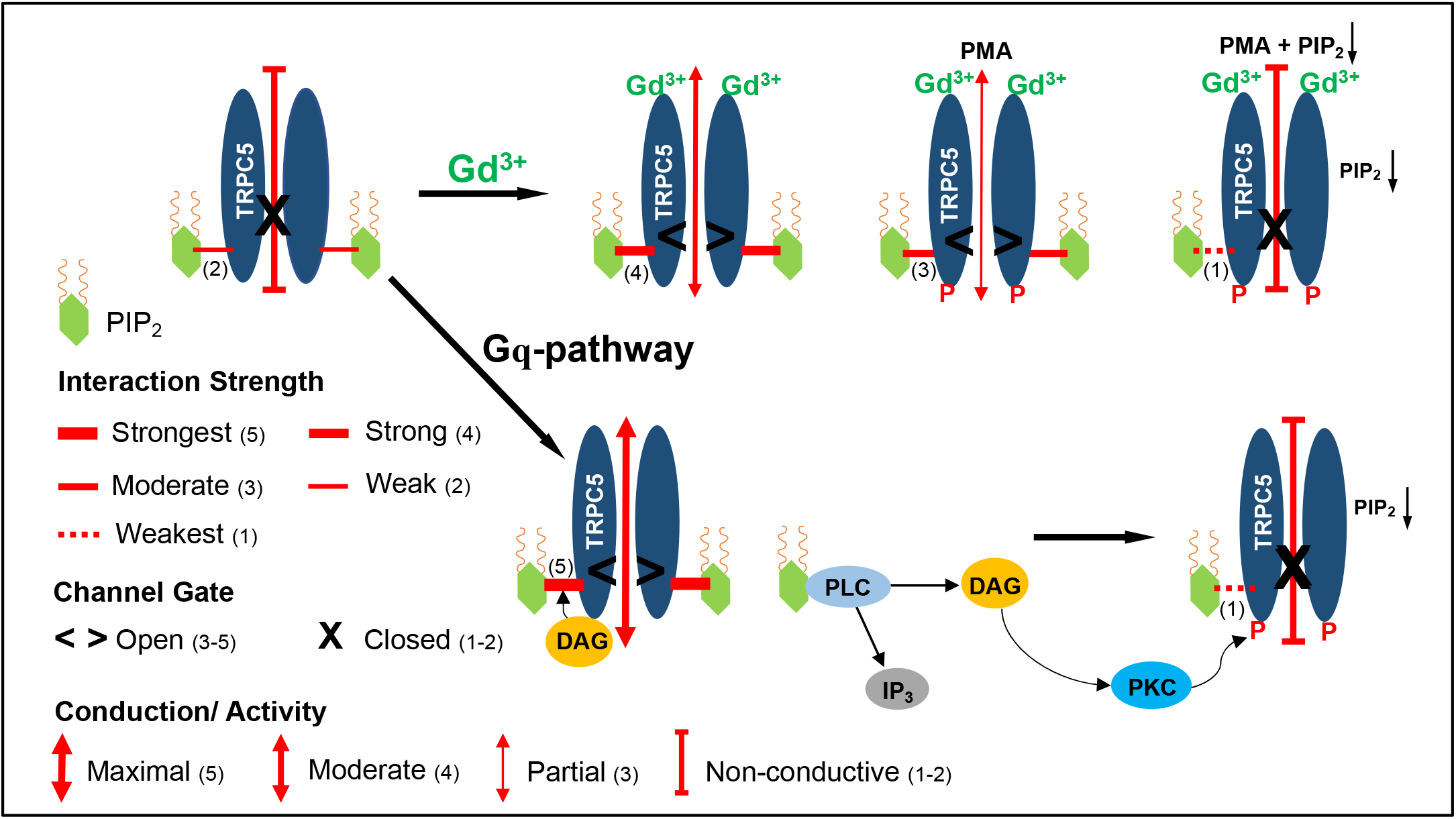
Cartoon model of the dependency of TRPC5 channel on PIP_2_ to account for stimulation and inhibition of channel activity by independent gating mechanisms. <u>Trivalent cation-mediated control of TRPC5 activity:</u> Trivalent cation activation mediated by Gd^3+^ allosterically strengthens channel interactions with PIP_2_, strongly enough to cause partial activation. PMA-treatment alone (PKC-mediated phosphorylation but not PIP_2_ depletion) weakens channel-PIP_2_ interaction strength and causes partial inhibition of channel currents. Similarly, depletion of intracellular PIP_2_ levels (using wortmannin or 5’-phosphatase) alone (PIP_2_ depletion but not PKC-mediated phosphorylation) does not strip the channel completely of its PIP_2_ causing partial inhibition of activity. The combination of PMA-treatment and PIP_2_ depletion strips the channel from its PIP_2_ severely enough to cause full inhibition. <u>Gq-mediated control of TRPC5 activity:</u> Upon Gq-receptor activation, PLC hydrolyzes PIP_2_ to IP_3_ and DAG. DAG allosterically enhances strongly channel interactions with PIP_2_ activating the channel maximally. PIP_2_ depletion (such as by dephosphorylation of PIP_2_) alone (without PKC-mediated phosphorylation as in T972A) causes partial inhibition. The ensuing DAG activation of PKC causes channel phosphorylation at T972, which allosterically weakens channel-PIP_2_ interactions enough, that adds up to the PIP_2_ depletion causing full inhibition of the current.

G_q/11_-mediated gating is more complex due to the fact that all four components, PIP_2_ depletion, DAG production and channel activation, DAG-mediated activation of PKC, and channel inhibition cannot be readily separated as with Gd^3+^ gating. The activating molecule DAG interacts with the intracellular side of the channel to strengthen channel-PIP_2_ interactions and yield maximal activation (strength 5/5), while protecting the channel from losing its PIP_2_ despite the ongoing PLC-meditated hydrolysis. In order for DAG to stimulate activity, the dominating inhibitory PKC phosphorylation needs to be abrogated. The obligate activation of PKC by the generated DAG molecules phosphorylates the channel at T972 weakening channel-PIP_2_ interactions (strength 1/5) and making the channel activation transient, as DAG-mediated activation is followed by complete inhibition of activity. Desensitization ensues as the hydrolyzed PIP_2_ needs to be resynthesized and protein phosphatases need to dephosphorylate the channel to render it activatable again by DAG. Our model proposes that even through Gd^3+^ and DAG use different pathways to activate TRPC5 channels they converge at the level of channel-PIP_2_ interactions which they control allosterically. Thus, in this model, it is PIP_2_ and its interactions with TRPC5 that should be deemed essential for channel activation and inhibition.

TRPC3/6/7 channels are highly sensitive to PLC-mediated PIP_2_ depletion which correlates with the spontaneous inhibition of DAG-activated currents observed (27). TRPC4/5 channels are uniquely regulated by C-terminal interactions with NHERF proteins, where in the NHERF-bound state the channel is DAG insensitive (2). Our proposed model suggests that PIP_2_ is significant for maximal channel activity and that a balance exists between PLC-mediated hydrolysis of PIP_2_ to make DAG which activates all of the TRPC channels and the remaining PIP_2_ molecules that are bound to the channel. When TRPC3-7 channels lose all of the bound PIP_2_, due to PKC-mediated phosphorylation that weakens channel-PIP_2_ interaction and the concurrent PIP_2_ hydrolysis by PLC enzymes, channel currents are fully inhibited. We speculate that an interplay exists between phosphorylated channel subunits and PIP_2_ bound subunits that lead to differences in activation and conduction in TRPC3-7 channels.

Previous studies performed to test the effect of PIP_2_ depletion using the PI3K and PI4K inhibitor Wortmannin showed that depleting PIP_2_ had no effect on CCh-mediated currents, whereas increasing PIP_2_ levels reduced the extent of CCh-mediated inhibition of TRPC5 channel currents (5, 30, 18). In an attempt to resolve these conflicting results, we incubated the cells with wortmannin to achieve the full effect and successfully deplete intracellular PIP_2_ levels and observed that wortmannin increased the rate of CCh-mediated inhibition and reduced the peak of Gd^3+^-activated currents. This conclusion was strengthened by the inability of increased PIP_2_ levels to prevent complete current inhibition during simultaneous PKC and 5’ phosphatase effect. This suggests that PKC-phosphorylation is dominant and, together with PIP_2_ depletion (PLC-mediated), results in irreversible inhibition of current that is unaffected by subsequent increases (diC_8_-PIP_2_) or decreases (by wortmannin) in PIP_2_ levels as previously shown (5,19, 28).

Fundamentally, the PIP_2_ dependency of several channels, including TRPC5, has been demonstrated in the inside-out patch configuration when, during patch excision, current rundown caused by a decrease in membrane PIP_2_ levels can be reversed by addition of PIP_2_ (8, 9). Trebak et. al established the role of an inhibitory factor that is associated with the channel in a PIP_2_ dependent manner which was later identified by Storch and colleagues to be the NHERF1/2 proteins (2, 5). However, the dependency of the affinity of the NHERF proteins for the channel on PIP_2_ levels remains to be explored. The underlying question to address going forward is whether channel-PIP_2_ binding physiologically accompanies NHERF protein binding and its inhibition of channel current.

Altogether we propose a model whereby channel-PIP_2_ interactions are important for TRPC5 channel activation as well as for maintenance of channel activity, assigning a critical functional role to PIP_2_ for this channel. We conclude that the dependence of channel activity on PIP_2_ levels may be a characteristic of all TRPC channels.

The broader relevance of this work is underscored by the fact that TRPC5 inhibitors are now being tested in Phase 2 studies in the clinic for the treatment of diseases caused by a leaky kidney filter, a direct consequence of the activation of TRPC5-mediated injury pathway in podocytes. A deeper understanding of the activation mechanisms of TRPC5 may enable the development of more selective and precise molecules to block TRPC5 channel activity in podocytes. Diseases driven by kidney filter damage, otherwise known as proteinuric or glomerular kidney diseases, account for the majority of the 850 million patients suffering from progressive kidney diseases worldwide. Therefore, our detailed studies can illuminate new therapeutic opportunities for targeted therapies for a group of diseases for which there is currently a great unmet need.

## Experimental Procedures

### Cell Culture

Human embryonic kidney (HEK293T) cells and Hippocampal HT-22 cells were acquired from ATCC (Manassas, VA) and maintained in DMEM (Sigma-Aldrich, Burlington, MA) supplemented with 100 units/mL penicillin, 100 μg/mL streptomycin and 10% (vol/vol) fetal bovine serum. All cells were held at 37 °C in a humified atmosphere with 5% CO_2_.

### Materials

OAG was purchased from Cayman Chemical (Ann Arbor, MI), diC8-PIP2 was from Echelon Biosciences (Salt Lake City, UT) and ML204 and AC1903 were received from Dr. Corey Hopkins’ Lab (University of Nebraska). PMA was purchased from LC labs (Woburn, MA), HEPES from Oakwood Chemical (Estill, SC). All other materials were purchased from Sigma-Aldrich (St. Louis, MO).

### Molecular Biology

Mouse TRPC5-GFP cDNA (<u>NM_009428.2</u>) constructs were used for whole-cell electrophysiology experiments. YFP was subcloned into the EGFP site/C-terminal end of the plasmid using respective restriction enzymes for TIRF experiments. Amino acid exchanges from Thr to Ala at position 972 in murine TRPC5 were introduced by site-directed mutagenesis using the QuikChange system (Stratagene). The cDNA constructs used in the present work were confirmed by Sangar sequencing (Macrogen, Cambridge, MA). CIBN-CAAX-GFP and CRY2-5’-ptase_OCRL_ were kind gifts from the DeCamilli lab (Yale, New Haven, CT). For TIRF experiments the constructs and were modified to remove existing fluorescent proteins.

### Electrophysiological Whole-Cell Measurements

HEK293T/HT-22 cells were seeded onto glass coverslips for whole-cell patch-clamp experiments. HEK293T cells were transfected with 2.5μg of mTRPC5-GFP, 2.5μg mTRPC5-T972A-GFP, 0.75μg CIBN-CAAX-GFP and 0.75μg CRY2-5’ptase-mCherry for the respective experiments using polyethylenimine (PEI). TIRF experiments were performed on cells transfected with 1 μg of YFP-mTRPC5 or YFP-TRPC5-T972A plus or minus, 0.75 μg CIBN-CAAX-GFP and 0.75 μg CRY2-5’ptase where indicated. All experiments were performed 18-24 hours after transfection. The standard pipette solution for patch-clamp contained, in mM: 140 CsCl, 2 EGTA, 10 HEPES, 0.2 Na_3_-GTP, and 2 MgCl_2_. The bath solution contained, in mM: 140 NaCl, 5 CsCl, 10 HEPES, 2 MgCl_2_, 2 CaCl_2_, 10 glucose. Both solutions were adjusted to pH7.4 adjusted with NaOH. The patch pipettes were made using a two-step-protocol (Sutter Instruments, Novato, CA) and had a resistance. between 5 to 8 MΩ. Once the whole-cell configuration was achieved, cells were clamped at a holding potential of −60 mV using a patch-clamp amplifier (Tecella, Foothill Ranch, CA), controlled by WinWCP software (University of Strathclyde, UK). Currents were studied using a voltage ramp stimulation from −100 mV to +100 mV applied every 1s. Data low pass filtered at 2 kHz, digitized at 10 kHz and analyzed using Clampfit (Molecular Devices, San Jose, CA). Liquid junction potentials were less than 3 MΩ and were not compensated for.

### Total internal reflection microscopy

Single fluorescent YFP-tagged mTRPC5 or mTRPC5-T972A channels were identified and studied at the surface of live HEK293T cells by TIRF microscopy, as described previously (1). Cells were seeded to #1.5 glass coverslips, transfected as described above and studied in the bath solution described above 18-24 hours after transfection. The evanescent wave for TIRF was established and calibrated to 100 nm using micro mirrors positioned below a high numerical-aperture apochromat objective (60x, 1.5 NA; Olympus, Waltham, MA) mounted on an RM21 microscope frame (Mad City Labs., Madison, WI) (2). YFP was excited by a 514-nm laser line (Coherent, Santa Clara, CA) and the emission was collected through a 540/ 30 nm bandpass filter (Chroma, Bellows Falls, VT) using an sCMOS back-illuminated camera (Teledyne Photometrics, Tucson, AZ) controlled by Micro-Manager freeware. Images were captured with a 200-ms exposure every 5-s. The surface density of single fluorescent channels was determined from 3-6 random 10 x 10 μm squares per cell (representing 100 x 100 pixels), and from 4-6 cells per group using the Analyze plugin in ImageJ. Intracellular application of OAG or PIP_2_ was accomplished by dialyzing the cytoplasm via a patch-pipette in whole-cell mode using the standard pipette solution described above. TIRF-patch mode was achieved using a micromanipulator mounted on the stage of the microscope to position the pipette, and development of whole-cell mode was monitored using a patch-clamp amplifier (Tecella, Foothill Ranch, CA) controlled by WinWCP software (University of Strathclyde, UK). To allow for full dialysis, cells were studied 200 s after whole-cell mode was established. Optogenetic dephosphorylation of PIP_2_ was performed in cells co-transfected with CRY2-5’-ptase_OCRL_ and CIBN-CAAX (see **Molecular Biology**) using 100-s illumination from a 445 nm laser line (Coherent, Santa Clara, CA).

### Statistical analysis

All statistical analyses were carried out using Graphpad Prism software. Results are presented as Mean ± SEM unless otherwise indicated. The comparisons were carried out using the Student’s t-test. P < 0.05 was considered statistically significant.

## AUTHOR INFORMATION

### Author Contributions

M.N and L.D.P performed research and analyzed data. M.N, L.D.P, A.G and D.E.L designed research. M.N and D.E.L wrote the paper.

## Acknowledgments

We thank the members of the D.E.L. laboratory. Takeharu Kawano for help with molecular biology and Heikki Vaananen for technical support. We are grateful to Yiming Zhou, Juan Lorenzo Pablo and James Pentikis for helpful discussions.

